# A fungal root endophyte functionally complements host immunity and mitigates natural immune variation in Arabidopsis

**DOI:** 10.64898/2025.12.02.691812

**Authors:** Natalie Peltz, Laura Armbruster, Christopher Stephens, Vivien Joisten-Rosenthal, Tim Thomsen, Alex W. Yan, Nick Dunken, Renan Granado, Stanislav Kopriva, Juliette de Meaux, Gregor Langen, Anna Koprivova, Björn Usadel, Alga Zuccaro

## Abstract

Beneficial root-associated microbes can enhance plant resilience by complementing aspects of host immunity. The fungal root endophyte *Serendipita indica* (*Si*) is known to promote plant growth and confer broad stress tolerance. To assess how natural host genetic variation influences *Si*-mediated protection, we screened 47 *Arabidopsis thaliana* accessions for susceptibility to the fungal pathogen *Bipolaris sorokiniana* (*Bs*) with and without *Si* colonization. All accessions benefited from *Si*, indicating that endophyte-mediated disease mitigation occurs broadly across diverse host genotypes. A focused comparison of two genetically and geographically proximate Swedish accessions, T510 and T530, which displayed the most divergent protection scores, revealed substantial differences in *Bs* susceptibility. Transcriptome profiling under bi- and tripartite colonization showed conserved defense responses in both accessions. *Bs* infection downregulated growth- and development-related genes, consistent with a growth–immunity trade-off, with T530 exhibiting higher *Bs* colonization and a stronger transcriptional response than T510. Co-colonization with *Si* effectively suppressed pathogen growth and disease symptoms in both accessions. Comparative genomic and transcriptomic analyses identified four immune receptor genes, including the TIR-NLR *ISI*, present in T510 but absent in T530. An *isi* T-DNA insertion mutant phenocopied the heightened *Bs* susceptibility of T530, confirming that *ISI* contributes to root immunity, while *Si*-mediated protection remained intact despite increased pathogen susceptibility. Together, these findings demonstrate that fungal endophytes can mitigate the functional consequences of natural immune variation and enhance the resilience of genetically diverse plant populations.

**Highlights:**

- *S. indica* confers broad protection against *B. sorokiniana* largely independent of host genotype or pathogen susceptibility.
- The TIR-NLR immune receptor *ISI* contributes to root immunity but is not essential for *S. indica*-mediated protection.
- Beneficial endophytes can mitigate natural immune variation effects and support overall plant health.

## Introduction

Plants host diverse communities of root-associated microbes, ranging from pathogens to mutualists. In geographically distinct *Arabidopsis thaliana* (*At*) populations, the composition of endophytic communities is strongly shaped by environmental factors such as soil characteristics and climate (Almario *et al*., 2017; Thiergart *et al*., 2020). Nonetheless, host genetic variation also plays a critical role in structuring the plant microbiome (Deng *et al*., 2021; Horton *et al*., 2014; Zhang *et al*., 2023). Genetic differences, particularly at loci involved in immune responses, hormone signalling, and cell wall integrity, can influence a plant’s ability to recruit specific microbial partners (Bulgarelli *et al*., 2012; Horton *et al*., 2014; Lundberg *et al*., 2012; Schlaeppi *et al*., 2014). Consequently, natural Arabidopsis accessions vary in their capacity to accommodate beneficial microbes and to defend against pathogens (Brachi *et al*., 2022). While genotype-dependent differences in bacterial communities have been widely explored, the functional interplay between host genetic variation and root-colonizing fungi remains comparatively understudied.

Fungi of the order Sebacinales are widespread mutualists with broad host ranges and are known to enhance plant growth and tolerance to both biotic and abiotic stress (Oberwinkler *et al*., 2013; Tedersoo *et al*., 2014; Weiß *et al*., 2016). Surveys of European *At* populations have shown that Sebacinales are significantly enriched in the rhizoplane of healthy roots, suggesting a conserved and functionally relevant endophytic association in natural settings (Mahdi *et al*., 2022). The Sebacinales fungus *Serendipita indica* (*Si*) has become a powerful model for dissecting interactions between beneficial and pathogenic microbes in the rhizosphere. In *At*, *Si* reduces colonization by the soil-borne pathogen *Bipolaris sorokiniana* (*Bs*) and mitigates *Bs*-induced disease symptoms (Eichfeld *et al*., 2024). As a hemibiotrophic fungus, *Bs* triggers strong defense responses in mono-associations but not when embedded in complex microbial communities (Mahdi *et al*. 2022), making it a useful tool for interrogating natural variation in host susceptibility and for assessing whether endophyte-mediated protection can compensate for differences in host immunity (Persson *et al*., 2009).

To investigate this, we screened 47 geographically and environmentally diverse natural *At* accessions for susceptibility to *Bs*, both with and without *Si*. Phenotypic and transcriptomic analyses of two genetically closely related Swedish accessions exhibiting the most divergent protection scores revealed substantial differences in *Bs* susceptibility and identified several immune components. Among these, the *Si*-induced TIR-NLR gene *ISI* (AT5G45240; Dunken *et al*., 2024) was experimentally confirmed to contribute to this variation. Importantly, *Si* conferred robust protection even in accessions lacking such immune factors, suggesting that endophyte-mediated disease mitigation can largely compensate for the functional impact of natural immune variation. These findings highlight a potential stabilizing role for mutualistic fungi in maintaining plant health across genetically diverse populations.

## Results

### *Si*-mediated protection against *Bs* is largely independent of host genotype and pathogen susceptibility

The beneficial root endophyte *Si* protects a wide range of host species from diverse phytopathogens (Bajaj *et al*., 2015; Li *et al*., 2022; Mensah *et al*., 2020; Narayan *et al*., 2017; Waller *et al*., 2005). However, the extent to which host genetic background influences this protective effect remains unclear. To investigate how natural genetic variation affects endophyte-mediated protection, we screened 45 European and two North American *At* accessions (Table S1) for *Bs*-induced disease symptoms in the presence and absence of *Si*. European accessions were prioritized because Sebacinales, including *Serendipita* spp., are recurrent in wild *At* populations across Europe (Mahdi *et al*., 2022), making them particularly relevant for evaluating naturally occurring plant–endophyte interactions. These accessions, drawn from the 1001 Genome Project (Alonso-Blanco *et al*., 2016; Weigel and Mott, 2009), span broad geographic and environmental diversity (Fig. 1A). Accessions were selected to maximize intra-continental genetic diversity (Fig. 1B). The two North American lines MNF-Che-2 and Tol-0 were included because North American *At* populations arose from multiple European introduction events and subsequently diversified locally (Shirsekar *et al*., 2021). Seedling health was quantified using pulse-amplitude modulation (PAM) fluorescence following inoculation with *Bs* alone or with both *Bs* and *Si*. A plant health index was integrated over time, and accession-specific protection scores were scaled to the Col-0 reference to account for biological variation across screening rounds (Fig. 1C).

**Figure 1:**
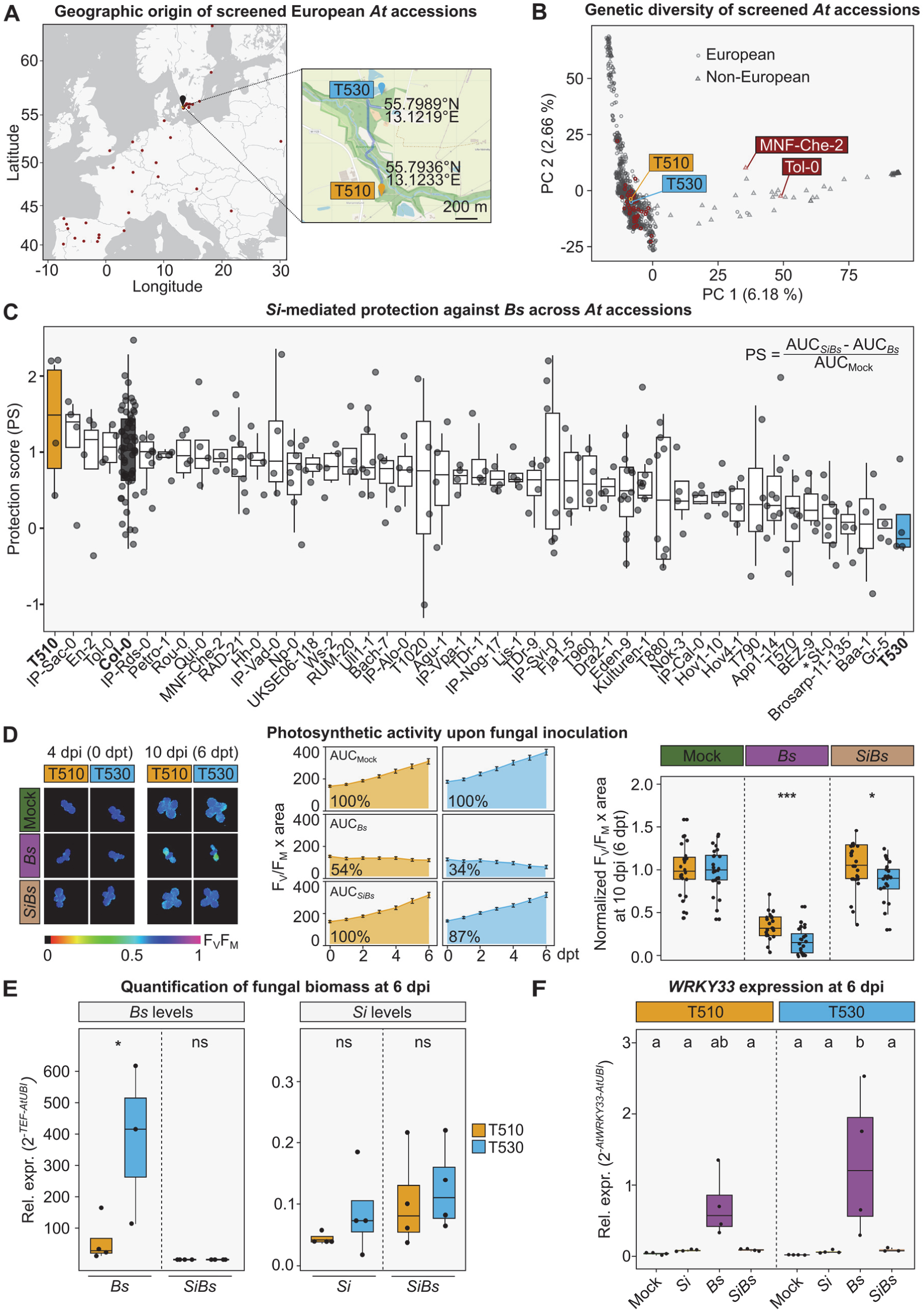
Genetic variation and *Si*-mediated protection against *Bs* in natural *At* accessions. **(A)** Geographic origin of screened European accessions. Map showing the geographical origin of natural European *At* accessions used in this study. Two North American accessions (MNF-Che-2 and Tol-0) were included as an outgroup but are not displayed. **(B) Genetic diversity among screened *At* accessions**. PCA plot based on MAF-filtered and LD-pruned SNP data from 1,135 *At* accessions included in the 1001 Genome Project (Alonso-Blanco *et al*., 2016). Screened accessions (highlighted in red) were selected to maximize representation across the spectrum of genetic diversity. **(C) *Si*-mediated protection against *Bs* across *At* accessions**. Seven-day-old seedlings from 47 accessions were treated with *Bs*, *SiBs*, or milliQ water (Mock) on 1/10 PNM Gelrite medium. At 7 dpi, seedlings were transferred to 48-well plates containing ddH2O, and plant health was monitored for 6 dpt using PAM fluorometry. Photosynthetic efficiency (FV/FM) was multiplied by projected plant area to obtain a plant health index. Health index values were integrated over time to calculate the AUC. Protection scores were derived from these AUCs and normalized to Col-0 to account for batch variation. The experiment was performed in five independent rounds with four plants per accession per treatment. Asterisk indicates statistically significant difference, observed only between St-0 and Col-0 (Kruskal-Wallis followed by Dunn’s test with Benjamini–Hochberg correction; p = 0.0478). **(D) Photosynthetic performance under pathogen stress and *Si*-mediated protection.** Left: Representative PAM fluorescence images for T510 and T530 at two time points under Mock, *Si* and *SiBs* treatments. Middle: Mean plant health over time with error bars representing standard error of the mean. Statistical significance of AUC values was assessed by two-way ANOVA followed by Tukey’s HSD *post-hoc* test. Right: Plant health at 10 dpi normalized to the Mock mean of each accession. Pairwise Wilcoxon rank-sum tests with Benjamini–Hochberg adjustment were used to assess statistical differences. n = 24. **(E) Quantification of fungal load across treatments and accessions.** Fungal load was quantified by qPCR using RNA extracted from root samples at 6 dpi. Fungal *TEF* gene expression was normalized to *AtUBI* (*UBQ5*, AT3G62250) and relative abundance was calculated using the 2^-ΔCt^ method. Statistical significance was evaluated using the Brunner–Munzel test with Benjamini–Hochberg correction per treatment between genotypes. n = 3-4. **(F) Defense gene expression in response to fungal colonization.** Expression levels of *WRKY33*, a central regulator of plant immunity against necrotrophic pathogens, in T510 and T530 at 6 dpi under Mock, *Bs*, *Si* and *SiBs* treatment. Statistical differences were assessed by one-way ANOVA followed by Tukey’s HSD *post-hoc* test. n = 3-4. Different letters indicate significant differences between groups (p < 0.05). Asterisks indicate p-values (* p<0.05, ** p<0.01, *** p<0.001).

All accessions showed positive protection scores, indicating that *Si* consistently mitigated *Bs*-induced disease symptoms across diverse host genotypes. Although protection scores ranged from 1.38 (T510) to 0.07 (T530), statistically significant differences were detected only between St-0 and Col-0. Notably, T510 and T530 represent opposite ends of the protection spectrum despite clustering closely in our genetic analyses and originating from the same local population in southern Sweden, where they were collected less than one kilometre apart (Fig. 1A and B). This combination of contrasting phenotypes and close genetic and geographic origin makes them a compelling pair for comparative analysis. When infected with *Bs*, both accessions exhibited pronounced disease symptoms, including root browning (Fig. S1A), reduced root length (Fig. S1B), and decreased plant health indices (Fig. 1D) relative to mock-inoculated controls. Consistent with the accession screen, co-inoculation with *Si* significantly reduced pathogen colonization levels and disease symptoms in both accessions (Fig. 1D and E). Pathogen load in mono-inoculated plants was approximately fourfold higher in T530 than in T510, whereas *Si* colonization levels were comparable between the two accessions at this time point (Fig. 1E). Both accessions showed induction of the immune marker gene WRKY33, a transcription factor known to activate camalexin pathway genes such as *CYP71A13* and *PAD3* (Petersen *et al*., 2008), upon *Bs* but not *Si* inoculation (Fig. 1F), with higher expression in T530 consistent with its elevated pathogen load. Co-inoculation with *Si* suppressed *Bs*-induced *WRKY33* expression in both accessions, in line with reduced *Bs* infection during tripartite interactions.

To further characterize the two accessions, we compared their growth under controlled laboratory conditions on axenic, nutrient-poor medium and in common-garden soil settings (Fig. S2A and B). Under laboratory conditions, T530 developed a slightly but significantly larger leaf surface area than T510 about two weeks after germination. However, in common-garden experiments extending to 56 days, no significant differences in growth or visible disease symptoms were observed between the accessions. These observations highlight that variation in growth and pathogen susceptibility measured under controlled laboratory conditions does not necessarily reflect plant performance in more natural environments, where interactions with resident microbial communities and variable abiotic factors, both of which can modulate growth–defense trade-offs, shape phenotypic outcomes.

Taken together, our findings (Fig. 1C–F) indicate that *Si*-mediated protection against *Bs* is robust across diverse *At* genotypes and is not readily explained by differences in *Si* colonization levels or by inherent differences in host susceptibility to *Bs*. Although the quantitative magnitude of protection differed among accessions, all lines exhibited a significant and consistently positive protective response, indicating that *Si*’s beneficial effect is largely genotype-independent. This interpretation aligns with previous observations that *Si* exerts much of its protective activity in the rhizosphere and rhizoplane, where it secretes antimicrobial effectors that inhibit *Bs* growth (Eichfeld *et al*., 2024).

### *Si* mitigates pathogen-induced stress responses and restores growth-related gene expression

To better understand the molecular basis underlying the differential susceptibility of T510 and T530 to *Bs* and to examine how *Si* modulates host responses, we performed transcriptome profiling of seedlings subjected to Mock, *Si*, *Bs*, and *Si*+*Bs* (*SiBs*) treatments. Roots were harvested three days post inoculation (dpi) and processed for RNA sequencing (RNA-seq) to capture early transcriptional responses to fungal colonization. To enable direct comparison of gene expression between T510 and T530, RNA-seq reads were aligned to the TAIR10 reference genome using Ensembl Plants release 58 gene annotations (sample-level mapping statistics in Table S2). Principal Component Analysis (PCA) of global transcriptional profiles revealed clear separation of samples by both host genotype and treatment (Fig. 2A). *Si*-treated samples clustered closest to the mock controls, indicating that *Si* alone induces only modest transcriptional changes at this early stage. By contrast, *Bs* treatment caused a pronounced shift in gene expression, reflecting a strong host response to pathogen infection. Samples co-inoculated with *Si* and *Bs* occupied an intermediate position in PCA space but clustered more closely with *Si*-treated samples than with *Bs* alone, indicating that *Si* attenuates the transcriptional changes induced by *Bs*. This transcriptional pattern is consistent with the observed reduction in *Bs* pathogen load.

**Figure 2:**
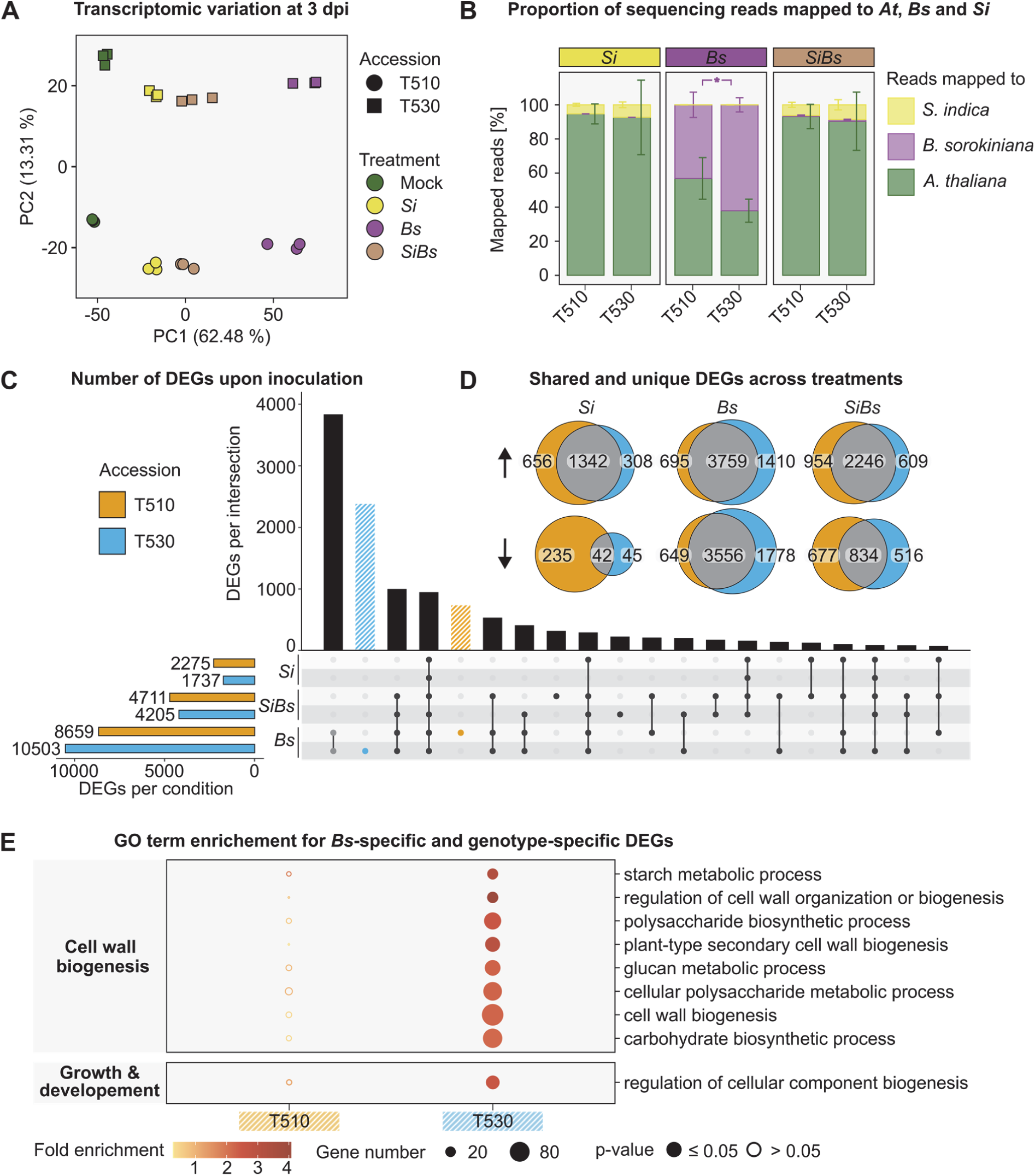
Transcriptional responses of Swedish *At* accessions to fungal colonization. **(A) Sample-level transcriptomic variation.** PCA plot based on normalized expression data depicting transcriptomic variation across genotypes (T510 and T530) and treatments (Mock, *Si*, *Bs*, *SiBs*). **(B) Mean proportion of sequencing reads mapping to *At*, *Si*, and *Bs*.** *Bs* fungal load was compared using a Student’s t-test (p = 0.0293). Error bars indicate standard deviation. n = 3. **(C) Upset plot showing the number of DEGs** (|log_2_FC| ≥ 1 and adjusted p-value <0.05). DEGs were identified in T510 and T530 in response to colonization by *Si*, *Bs* and *SiBs* relative to Mock (n = 3). Up and downregulated DEGs are included. **(D) Venn diagram illustrating the overlap of DEGs across fungal treatments.** Up- and downregulated DEGs were separated for this analysis. n = 3. **(E) GO enrichment analysis of DEGs uniquely responsive to *Bs* in T510 or T530.** Analysis was conducted on sets of DEGs highlighted in (C). Dot color indicates fold enrichment, color fill denotes FDR significance, and dot size reflects the number of genes associated with each GO term.

To estimate the relative abundance of each organism within the samples, RNA-seq reads were mapped to the reference genomes of *At* (TAIR10, Table S2), *Si* (Pirin1; Zuccaro *et al*. (2011), Table S3) and *Bs* (Cocsa1; Ohm *et al*. (2012) and Condon *et al*. (2013), Table S4). Reads mapping ambiguously to more than one genome were excluded. Based on the proportion of uniquely mapped reads, *Si* colonization did not differ significantly between T510 and T530 under any treatment (Fig. 2B). Consistent with earlier observations (Fig. 1E), *Bs* colonization varied between accessions in the absence of *Si*, with T530 exhibiting a higher mean proportion of *Bs*-derived reads (62%) than T510 (43%). In both accessions, co-inoculation with *Si* markedly suppressed *Bs* colonization, reducing its relative abundance to below 1%, while *Si* colonization remained stable under co-treatment conditions, comprising 5–9% of total reads in both *Si*- and *SiBs*-treated samples. To examine transcriptional plant responses to fungal treatments, we identified differentially expressed genes (DEGs; |log₂FC| ≥ 1, adj. p ≤ 0.05) by comparing each treatment to its respective mock control (Fig. 2C; full results in Table S5). *Bs* elicited the strongest transcriptional response in both accessions, with T530 displaying more DEGs (10,503) than T510 (8,659), likely reflecting its greater *Bs* colonization. Co-inoculation with *Si* substantially dampened this response, reducing the number of DEGs by nearly 50% in both accessions (to 4,711 in T510 and 4,205 in T530). By contrast, *Si* treatment alone induced comparatively modest transcriptional changes (2,275 DEGs in T510 and 1,737 in T530). To further characterize these responses, DEGs were partitioned into up- and downregulated sets and analyzed by accession (Fig. 2D). Although a substantial fraction of genes was commonly regulated across accessions in response to microbial treatments, T510 exhibited a stronger accession-specific response under both *Si* and *SiBs* conditions, evident in both upregulated and downregulated gene sets. In contrast, T530’s accession-specific signature was predominantly associated with *Bs* treatment. These findings support the presence of a conserved core transcriptional response to fungal colonization, modulated by accession-specific programs reflecting differential sensitivities to individual microbial cues (Fig. 2E).

Gene ontology (GO) enrichment analysis (Fig. S3, Table S6) revealed that both accessions responded to *Bs* treatment with strong induction of immune-related processes. These signatures included enrichment of classical defense-associated phytohormone pathways involving salicylic acid (GO:0009751), ethylene (GO:0009723), and jasmonic acid (GO:0071395), as well as secondary metabolism. A hallmark of the *Bs* response was the activation of cell death–related processes (GO:0008219), which were not observed under *Si* or *SiBs* treatments. *Si* alone induced only mild changes in defense- or stress-associated categories. Strikingly, *Bs* infection caused strong downregulation of genes associated with plant growth and developmental processes, including the mitotic (GO:0000278) and meiotic (GO:0051321) cell cycles, epidermal cell differentiation (GO:0090627), and cell maturation (GO:0048469), reflecting a canonical growth–defense trade-off. Co-treatment with *Si* largely restored these processes and mitigated *Bs*-induced stress pathways associated with reactive oxygen species (GO:0000302) and nutrient deprivation (GO:0031667), further supporting a protective role of *Si* in modulating defense responses, stress and growth inhibition triggered by *Bs* infection.

Given the differential susceptibility of T510 and T530 to *Bs* colonization, we examined accession-specific transcriptional responses to the pathogen (Fig. 2C,E). In *Bs*-treated T510, no significantly enriched GO terms were detected among uniquely differentially expressed genes. In contrast, the accession-specific response of T530 showed significant enrichment of GO terms related to cell wall biogenesis (GO:0042546), cell wall organization (GO:0071554), and polysaccharide biosynthesis (GO:0000271), with a particular emphasis on glucan biosynthesis (GO:0044042). These findings indicate that the accession-specific response of T530 to *Bs* involves transcriptional changes in pathways related to host cell wall remodelling and structural reinforcement, although it remains unclear whether these changes contribute to, or arise from, its heightened susceptibility.

### T510 and T530 display presence–absence variation (PAV) in immune-related genes

To investigate whether structural variation contributes to the differential *Bs* colonization patterns in T510 and T530, we leveraged the transcriptome data to screen for potential gene presence–absence variation between the two accessions. For this purpose, we examined RNA-seq datasets from T510 and T530, and additionally included Col-0 as a well-annotated reference accession to help distinguish true PAV from low-expression artifacts. Genes showing very low expression (one or fewer average counts) in one genotype but robust expression (ten or more average counts) in the other two across three biological replicates and all four treatments (Mock, *Si*, *Bs*, *SiBs*) were considered candidate presence–absence variants. This analysis identified 58 candidate genes potentially absent in T510 and 48 candidate genes potentially absent in T530 (Fig. 3A). Most of these candidates were protein-coding (45/58 in T510 and 37/48 in T530), with more than one-third lacking annotated Pfam domains. Among those with predicted functions, the most frequently represented domain was the TIR domain, present in ten genes, nine of which appeared absent in T530. These findings are consistent with previous reports of extensive presence–absence variation among immunity-related genes in *At* (Bush *et al*., 2014; Van de Weyer *et al*., 2019).

**Figure 3:**
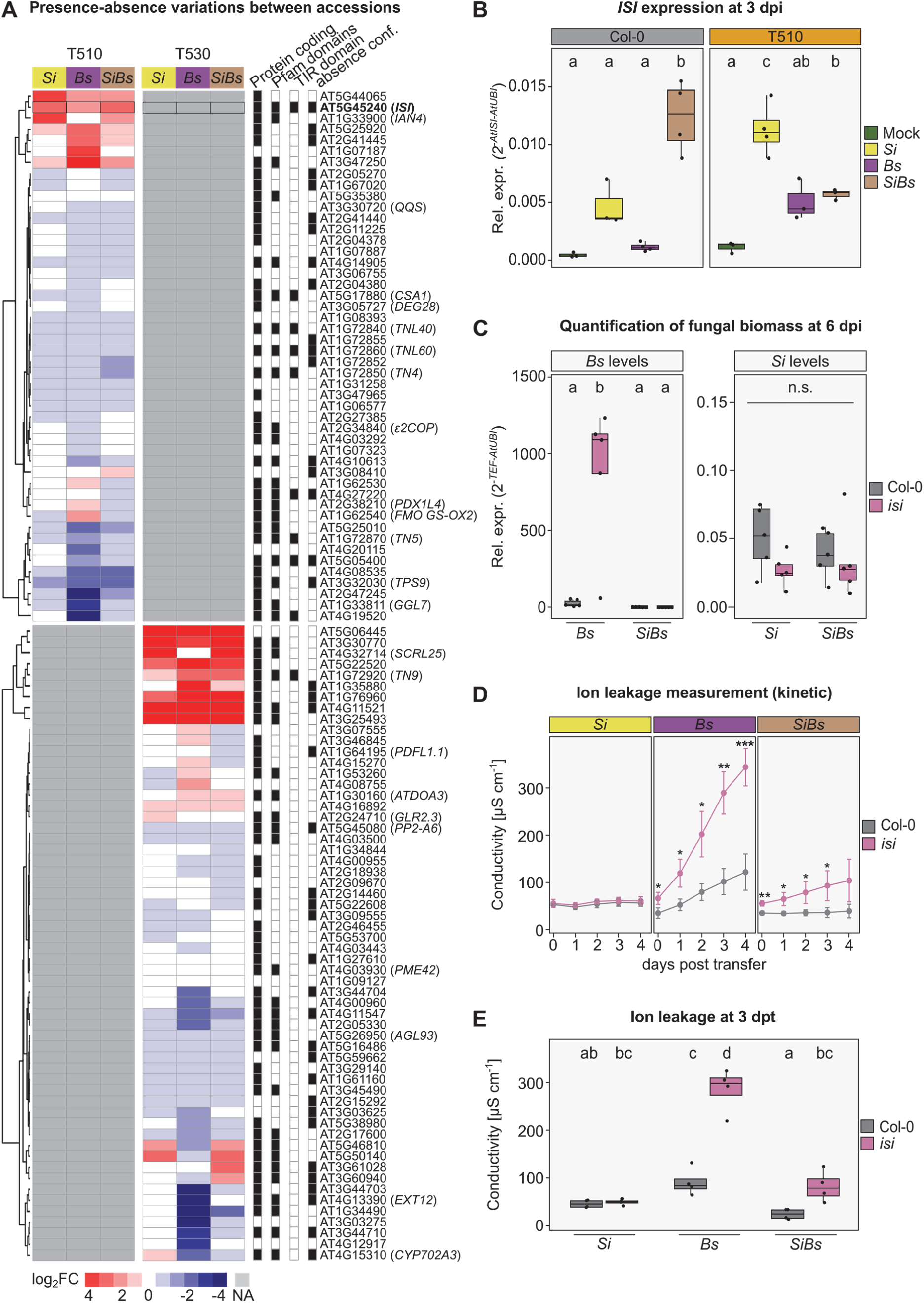
The absence of the TIR-NLR *ISI* enhances susceptibility to *Bs* but does not impact *Si*-mediated protection. **(A) Heatmap depicting expression of genes subject to presence-absence variation in T510 and T530 after *Si*, *Bs* and *SiBs* treatment as log2FC with respect to Mock.** To identify genes expressed in only one of the two Swedish accessions, the RNA-seq data of Col-0, T510 and T530 were screened for genes with one or fewer average counts in one genotype and ten or more average counts in the other two genotypes over three biological replicates and four treatments (Mock, *Si*, *Bs*, *SiBs*). Prediction of Pfam domains using InterProScan (https://www.ebi.ac.uk/interpro/) was used to identify TIR domain-containing proteins. Of these, only one (AT5G45240, *ISI*) was strongly induced after *Si*, *Bs* and *SiBs* treatment in T510. Predicted protein structure is represented to the left of gene identifiers, and the rightmost column shows true presence-absence based on genome analysis. **(B) *ISI* gene expression**. Gene expression was quantified by qPCR using RNA extracted from root samples at 3 dpi (n = 3-4). *ISI* gene expression was normalized to *UBI* gene expression as a reference, and relative abundance was calculated using the 2^-ΔCt^ method. Statistical significance was assessed separately for each genotype using one-way ANOVA followed by Tukey’s HSD *post-hoc* test. **(C) *Bs* colonization and *Si*-mediated protection in *isi* T-DNA line.** Fungal load was quantified by qPCR using RNA extracted from root samples at 6 dpi (n = 5). *Bs* TEF gene expression was normalized to *AtUBI* as a reference, and relative abundance was calculated using the 2^-ΔCt^ method. Statistical differences were assessed by two-way ANOVA followed by Tukey’s HSD *post-hoc* test. **(D) Time course of ion leakage upon *Bs* infection and *Si*-mediated protection in the *isi* mutant.** Seven-day-old plants were treated with *Si* mycelium, *Bs* spores or a combination of both. After five days, the plants were transferred to wells filled with milliQ water. Conductivity (µS x cm^-1^) was measured from 0 to 4 dpt with two technical replicates per biological replicate (n = 4). Data are displayed as average ± standard error of the mean. A Student’s t-test was performed to compare Col-0 and the *isi*-mutant for each condition and timepoint separately (asterisks indicate significance level), * p < 0.05, ** p < 0.01, *** p < 0.001). **(E) Ion leakage in inoculated *isi* seedlings**. Ion leakage was measured at 3 dpt for Col-0 and *isi* mutants under *Si*, *Bs*, and *SiBs* treatments, as shown in (D). Conductivity values were log-transformed and analysed using a linear model (LM), followed by ANOVA and Tukey’s *post-hoc* test. Statistically significant differences (p < 0.05) are indicated by distinct letters.

### Deletion of TIR-NLR *ISI* in Col-0 phenocopies the increased *Bs* susceptibility of T530

Notably, only one of the nine TIR-domain–containing genes present in T510 but absent in T530, *AtISI*, was strongly induced by all fungal treatments (Fig. 3A,B). *ISI* encodes a TIR-NLR protein recently identified as a modulator of *Si*-induced, regulated cell death in roots at later colonization stages (7–10 dpi; Dunken *et al*., 2024). To evaluate whether *ISI* contributes to the differential *Bs* susceptibility phenotype, we quantified fungal colonization in a Col-0 *isi* T-DNA insertion knockout (SALK_034517) following *Si*, *Bs*, and *SiBs* treatments at the same early time point used for transcriptomics (3 dpi). Similar to T530, the *isi* mutant displayed pronounced hyper-susceptibility to *Bs*, which was strongly suppressed by co-inoculation with *Si* (Fig. 3C). Consistently, *SiBs* treatment attenuated *Bs*-induced disease symptoms in *isi* seedlings, as evidenced by reduced ion leakage (Fig. 3D,E), a proxy for loss of membrane integrity associated with cell death. Despite its functional relevance, *ISI* is absent in several accessions within our panel (Table S7), highlighting substantial natural variation at this immune-related locus.

To examine whether *ISI* contributes more broadly to microbial colonization beyond fungal interactions, we assessed colonization levels in *isi* seedlings three days after inoculation with the bacterial pathogen *Burkholderia glumae* PG1 (*Bg*), a motile, soil-borne bacterium. *Bg* is the causal agent of bacterial panicle blight in rice (Ham *et al*., 2011) and causes bacterial wilt in various crops (Kim *et al*., 2014). In *At*, *Bg* elicits strong immune responses and growth inhibition, including induction of camalexin production (Koprivova *et al*., 2019). Three days after mono-inoculation with *Bg*, no significant differences in bacterial load were observed between Col-0 and the *isi* mutant, indicating that *ISI* is not required to restrict *Bg* colonization (Fig. 4A). As previously observed for *Bs*, co-inoculation with *Si* markedly reduced *Bg* colonization in both genotypes, suggesting that *Si* confers broad-spectrum protection against both fungal and bacterial pathogens.

**Figure 4:**
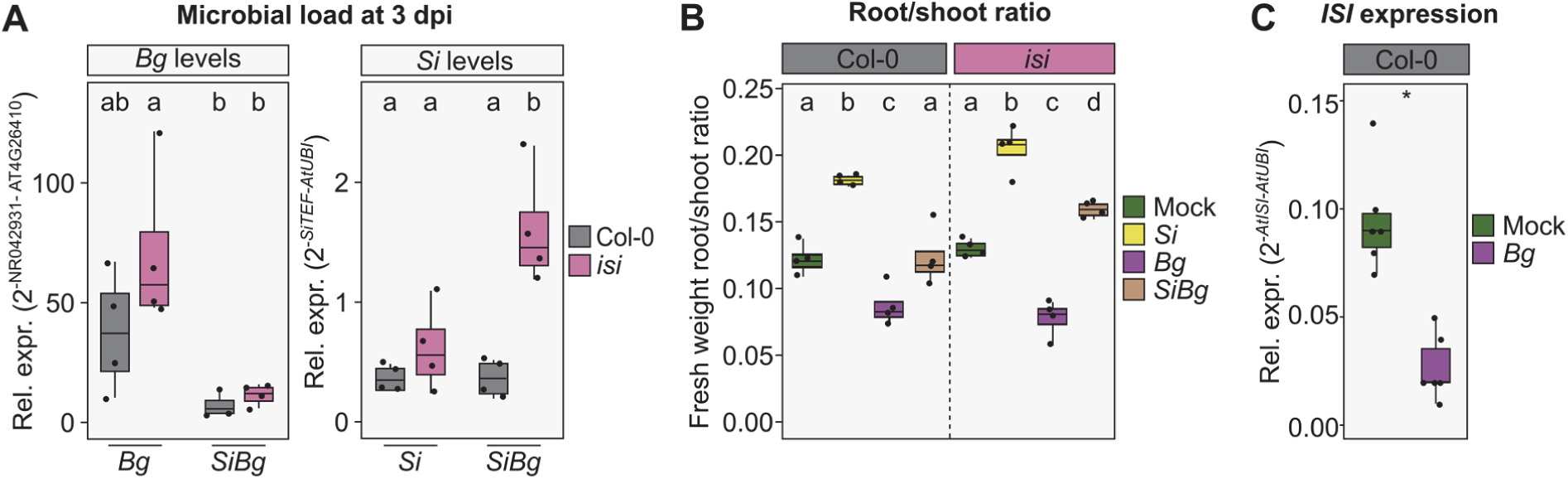
*Si* protects plants from the bacterial pathogen *Bg* independent of the presence of the immune receptor *ISI*. **(A) Microbial colonization of *isi* seedlings at 3 dpi.** *Bg* NR042931 expression was normalized to *AtRHIP1* (AT4G26410) as established in Ross and Somssich (2016) (left) and *SiTEF* gene expression was normalized to *AtUBI* as a reference (right). Relative abundance was calculated using the 2^-ΔCt^ method. Data are shown per genotype and treatment (n = 4). Two-way ANOVA followed by Tukey’s *post-hoc* test was applied. Statistically significant differences (p < 0.05) are indicated by different letters. **(B) Fresh weight of plant root vs. shoot tissue upon microbial inoculation at 3 dpi (n = 4).** Two-way ANOVA followed by Tukey’s HSD was performed. **(C) *ISI* gene expression upon *Bg* treatment**. Gene expression was quantified by qPCR using RNA extracted from root samples at 3 dpi (n = 6). *ISI* gene expression was normalized to *UBI* gene expression as a reference, and relative abundance was calculated using the 2^-ΔCt^ method. Statistical significance was assessed via a Student’s t-test (p < 0.05) and significant differences are indicated by asterisks.

While *Si* colonization remained stable in Col-0 during *Bg* co-inoculation, the *isi* mutant exhibited a pronounced increase in *Si* colonization (Fig. 4A), a pattern not observed during *Bs* co-inoculation (Fig. 3C). Previous work showed that *ISI* restricts *Si* proliferation during later colonization stages, with loss of *ISI* leading to elevated cell death and *Si* hypercolonization (Dunken *et al*., 2024). The elevated *Si* levels in *Bg*-treated *isi* seedlings may reflect an accelerated or enhanced colonization dynamic under these conditions. This suggests *Bg*-induced changes to the host environment or immune responses that facilitate more rapid or extensive *Si* colonization when *ISI*-mediated restriction is absent. Differences in *Si* colonization between *Bg* and *Bs* co-infections likely reflect distinct pathogen interactions, metabolic effects, or immune modulation pathways triggered by bacterial versus fungal pathogens.

To assess whether bacterial infection triggers additional *ISI*-dependent host phenotypes, we monitored plant growth after inoculation. As the mesh-based setup used for bacterial inoculation prevented precise separation of individual roots and shoots, we used the bulk root-to-shoot ratio as a normalized and sensitive indicator of early growth perturbations. Mono-inoculation with *Bg*, but not *Si*, significantly reduced the root-shoot biomass ratio in both genotypes (Fig. 4B). Root growth arrest is a common response to microbial signals, including microbial-associated molecular patterns such as flg22, plant hormones, and volatile compounds (Dini-Andreote *et al*., 2025). Consistent with prior reports that *Si* suppresses flg22-induced growth inhibition (Jacobs *et al*., 2011), *Si* co-treatment mitigated the negative impact of *Bg* on root–shoot allocation independent of *ISI* status. This is further supported by gene expression data showing that *Bg*, unlike *Si*, did not induce *ISI* at this stage and instead significantly suppressed its expression (Fig. 4C), indicating that *ISI* is not engaged during bacterial colonization at this stage.

### Long-read genome assemblies reveal substantial immune-gene structural variation between T510 and T530

To validate structural variation in immunity-related genes, including *ISI*, in the accessions T510 and T530, we generated high-quality genome assemblies using Oxford Nanopore long-read sequencing. The resulting assemblies showed chromosome-scale contiguity and high completeness (Table S8). Whole-genome alignments using SyRI revealed substantial overall synteny between both accessions and the Col-0 reference genome, with several small inversions distributed across all chromosomes and a prominent inversion on chromosome 4 (Fig. S4). Genome annotation was performed by lifting over the TAIR10 reference annotation, supplemented with a *de novo* gene annotation generated by Helixer (Holst *et al*., 2023; Stiehler *et al*., 2021) and supported by RNA-seq based evidence.

We focused our analysis on the candidate genes previously identified as putatively absent in one accession but not the other based on their contrasting expression patterns (Fig. 3A). In T510, 58 candidate genes were examined (Table S9). Of these, 40 % (23/58) were confirmed to be truly absent at the genome level. Most non–protein-coding candidates were present but simply not expressed (only 4 of 13 were truly absent), whereas nearly half of the protein-coding candidates were absent (19/45). In nine cases, the best genomic alignments corresponded to paralogous loci but still exhibited substantial sequence identity (>60 %) to the Col-0 reference, supporting classification as present. One exceptional case (AT5G22520) showed only ∼50% identity to the reference but was still classified as present due to the absence of more similar matches. In T530, 35 % of the 48 candidate genes (17/48) were confirmed absent (Table S10). Among non–protein-coding candidates, 3 of 11 were truly absent; among protein-coding genes, 14 of 37 were absent. For several genes classified as present, sequence identity to Col-0 was relatively low (∼54–60%), indicating divergence rather than true gene loss. In six cases, alternative paralogs showed higher similarity to the candidate gene than the Col-0 reference, yet the gene still exhibited ≥60% identity within the syntenic region and was therefore considered present. Conversely, genes were classified as absent if their closest alternative match shared <60% identity (6 of 14) or if no homologous sequence was detected at all (8 of 14). Notably, *ISI* belonged to the latter group. In addition to *ISI*, we confirmed the absence of three further TIR-domain–containing genes in T530.

Hence, we examined the local genomic architecture surrounding the *ISI* locus in greater detail (Fig. 5, right panel). In T510, the corresponding region on Chromosome 5 is highly syntenic with Col-0, consistent with the presence of *ISI*. In contrast, T530 displays a rearranged structure, including a translocation involving *AT5G45230* and a clear deletion of *ISI*. The T530 locus contains several additional genes not present in Col-0 or T510, including *CHR5G3961* and *CHR5G3955*, both nearly identical to predicted TIR domain–containing proteins in other accessions (KAL9813235.1 and KAG7604990.1, respectively), as well as *CHR5G3964*, which shows sequence similarity to an LRR protein (KAG7532627.1). These rearrangements show local cluster variation and remodelling of immune-related loci in T530, consistent with high variability in this region.

**Figure 5:**
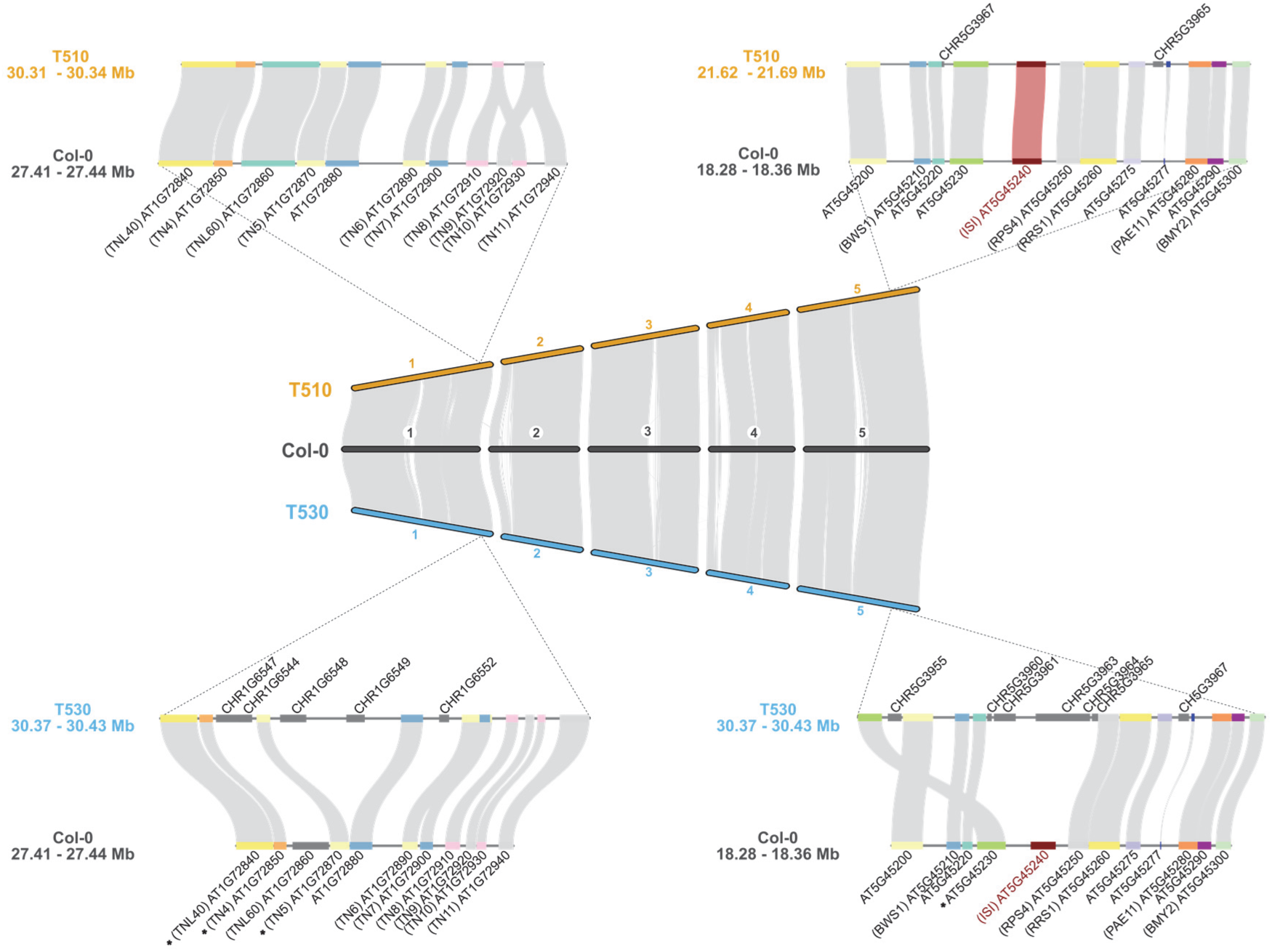
Macro- and microsynteny analysis of T510 and T530 compared to the TAIR10 Col-0 reference genome of *A. thaliana*. The central panel shows the macrosynteny relationships across all five chromosomes between the accessions. Syntenic gene regions are depicted as grey connecting lines between genomes. Gene codes marked with an asterisk (*) indicate syntenic blocks that exhibit less than 60% sequence identity with the corresponding genes in T530. The left panel presents an enlarged view of the macrosynteny on Chromosome 1, highlighting the region spanning genes AT1G72840–AT1G72940 in comparison to Col-0. The right panel shows a detailed microsynteny view of Chromosome 5 covering the interval AT5G45200–AT5G45300, which includes the *ISI* gene (AT5G45240, marked in red). While the *ISI* gene and its surrounding syntenic block are conserved in T510, this region lacks a corresponding syntenic block in T530.

We also examined a region on Chromosome 1 enriched in TIR–NBS–encoding genes that contains another TIR-NLR gene absent in T530 (Fig. 5, left panel). In T510, the interval spanning 30.31–30.34 Mb shows strong synteny with the Col-0 region extending from *TNL40* (*AT1G72840*) to *TN7* (*AT1G72900*). Notably, both Col-0 and T530 contain a duplicated segment spanning *TN8* (*AT1G72910*) through *TN11* (*AT1G72940*), whereas in T510 this region is present only as single-copy representations of *TN8/TN10* and *TN9/TN11*. In addition, two extra genes are located between *TN5* (*AT1G72870*) and the *surE*-like phosphatase locus (*AT1G72880*) in T530. One of these, *CHR1G6549*, shows perfect protein-level alignment to a TIR domain–containing gene from *Arabidopsis suecica* (KAG7659361.1; Chen, unpublished), while the second gene lacks detectable homology to any known protein.

Together, these analyses confirm substantial structural variation between T510 and T530, particularly within immune-gene clusters, providing a genomic context that can help explain differences in their immune capacities.

## Discussion

Plants and their associated microbiomes function as integrated ecological units, or holobionts, in which microbial partners extend host capacities in growth, nutrient acquisition, and tolerance to biotic and abiotic stresses (Berg *et al*., 2024; Hawkes *et al*., 2021; Mesny *et al*., 2023). This integrated cooperation is particularly relevant in the context of plant immunity, where beneficial microbes can augment host disease resistance and complement innate immune pathways (Vannier *et al*., 2019). Here, we show that the beneficial root endophyte *Si* protects Arabidopsis seedlings from the fungal pathogen *Bs* across a diverse panel of European accessions. Quantitative differences between accessions were small relative to the overall protective effect, indicating that *Si* enhances disease suppression across a broad range of host genetic backgrounds. The widespread association of Sebacinales with wild *At* ecotypes suggests that mutualistic fungi may act as ecological stabilizers of plant performance across diverse immunogenetic backgrounds and heterogeneous environments (Mahdi *et al*., 2022).

Genome-wide association studies (GWAS) are a powerful approach for identifying genetic determinants of quantitative traits, but their resolution is inherently limited in small datasets. Moreover, the complexity of plant-microbe interactions poses additional challenges for GWAS, particularly when trait variation is driven by PAV, transcriptional plasticity, or microbe- and context-dependent effects (Tam *et al*., 2019). We therefore complemented population-level approaches with a targeted comparison of two genetically similar Swedish accessions, T510 and T530, to dissect phenotypic variation. Although both accessions were protected by the beneficial fungus, T530 displayed greater susceptibility to the fungal pathogen. While the host genetic background can broadly influence microbiome assembly, discrete immune loci often exert disproportionate effects on specific plant–microbe interactions (Janse van Rensburg *et al*., 2025). In *At*, immunity relies on a diverse repertoire of intracellular NLR receptors, many of which belong to the TIR-NLR subclass that trigger strong defense responses upon effector recognition (Ngou *et al*., 2021). The recently characterized TIR-NLR *ISI* modulates fungal colonization and cell death in roots and is induced by *Si* and *Bs*, yet is absent in a substantial fraction (∼76 %) of natural *At* accessions, including T530. Loss of *ISI* in Col-0 is sufficient to phenocopy the heightened *Bs* susceptibility observed in T530, supporting its contribution to root immunity. Nonetheless, additional genetic factors almost certainly contribute to the susceptibility differences between T510 and T530, including variation in other immune receptors, basal hormone signalling, or accession-specific transcriptional regulation of defense.

Comparative genomic analysis revealed that in T530 the *ISI* locus is replaced by a cluster of NLR- and TIR domain–containing genes not found in Col-0 or T510, indicating local cluster variation and structural remodeling at this highly variable immune-gene region (Lee and Chae, 2020; Saile *et al*., 2020). Despite the presence of these additional predicted genes, T530 remains more susceptible to *Bs*, suggesting that they do not compensate for *ISI* loss under the tested conditions. Together with the *isi* knockout phenotype, these findings indicate that *ISI* contributes to fungal resistance in the roots. The lack of *ISI* induction, and its suppression, during *Bg* infection, along with the absence of detectable effects on *Bg* colonization, suggests that *ISI* function is engaged in a microbe-specific and context-dependent manner. Yet, *Si* still conferred strong protection against *Bg* in both Col-0 and the *isi* mutant, demonstrating that *Si*-mediated mitigation extends beyond a single pathogen class and is largely independent of *ISI*. Whether the remodelled TIR-NLR cluster in T530 provides alternative recognition specificities or functions against other pathogens remains unknown. Similar rearrangements in NLR clusters across *At* and other Brassicaceae are known to modulate effector recognition and immune output (Barragan *et al*., 2019; Goritschnig *et al*., 2016), suggesting that this region may retain functional significance despite *ISI* loss.

In holobiont systems, the selective pressure to maintain costly immune receptors may be reduced when beneficial microbes buffer the fitness consequences of receptor loss (Mesny *et al*., 2023; Schneider, 2021). Our results support this principle: *Si* provides robust protection even in accessions lacking *ISI*, potentially contributing to a context in which immune-gene PAV can accumulate without immediate fitness penalties in natural settings. This interplay between genomic rearrangement, microbial buffering, and functional immunity likely contributes to the extensive natural diversity of NLR gene content across *At* populations.

Among the TIR domain–containing genes with PAV between T510 and T530, *ISI* was the only one strongly induced by fungal treatments. However, other PAV genes may still contribute to immunity in ways not captured here. Their activity could be confined to specific cell types (Tang *et al*., 2023), developmental stages (Lüdke *et al*., 2025), or tissues not sampled, or they may respond to distinct microbial taxa. Alternatively, some may act as constitutive surveillance components that do not require transcriptional induction. Further functional studies will be required to clarify their roles and to determine whether they contribute to pathogen specificity or fine-tuning of immune signalling.

Despite the overall conservation of genome structure between T510 and T530, the two examined immune gene clusters stood out as regions of structural rearrangement. This is consistent with the view that NLR clusters act as evolutionary hotspots subject to recurrent birth–death dynamics and recombination processes that continuously reshape immune repertoires in response to fluctuating pathogen pressures (Michelmore and Meyers, 1998). Activation of TIR-NLRs often carries fitness costs (Brown and Rant, 2013; Burdon and Thrall, 2003), and the resulting selection pressures lead to extensive PAV and regulatory variation (Bush *et al*., 2014; Van de Weyer *et al*., 2019). These patterns reflect evolutionary trade-offs between defense and microbial compatibility in the rhizosphere where beneficial and pathogenic microbes coexist (Haney *et al*., 2015; Runge *et al*., 2023). Notably, T530 developed larger leaf surface area than T510 under controlled laboratory conditions but not in common-garden experiments, where neither accession showed overt disease symptoms. This suggests that simplified laboratory environments can amplify the apparent fitness costs of immune-gene variation, whereas natural or soil-like conditions buffer such effects (Giolai and Laine, 2024). Recent work supports this environment-dependent masking of immune trade-offs (Lundberg *et al*., 2025). Our findings extend this concept by showing that even a single beneficial microbe can mask susceptibility differences in controlled settings, highlighting microbial buffering as an additional mechanism that mitigates the fitness impact of immune-gene variation.

In summary, our findings support the hypothesis that beneficial microbes like *Si* can mitigate the consequences of immune gene loss and facilitate the persistence of allelic polymorphisms by functionally compensating for gaps in host immunity (Fig. 6). This aligns with emerging views that microbiome members can buffer the functional outcomes of genetic deficiencies and thereby influence host evolutionary trajectories (Henry *et al*., 2021). Beneficial microbes protect their hosts through multiple mechanisms, including immune priming and transcriptional modulation (Stein *et al*., 2008; Molitor & Kogel 2009), secretion of antimicrobial effectors (Chen *et al*., 2018b; Eichfeld *et al*., 2024), competitive exclusion of pathogens (Zelezniak *et al*., 2015), and mycoparasitism (Parratt and Laine, 2018). Previous work linked *Si*-mediated protection to direct antagonistic activity between *Si* and *Bs* in the rhizosphere (Eichfeld *et al*., 2024). Our findings support this model by showing that *Si*-mediated protection is effective across diverse host genotypes. This microbial buffering likely reduces selective pressure to maintain costly immune receptors, reminiscent of the “Black Queen” hypothesis in which functions become outsourced to microbial partners (Morris, 2015). Over evolutionary timescales, such outsourcing may reshape host immune architectures by favouring the loss of energetically costly receptors and those that risk inappropriate activation or autoimmunity, while maintaining phenotypic robustness in plant holobionts that include fungal endophytes. This further suggests that effective pathogenesis requires adaptation to the holobiont as a whole, meaning that pathogens must circumvent not only host-encoded immunity but also the additional protective layer contributed by commensal and mutualistic microbes.

**Figure 6:**
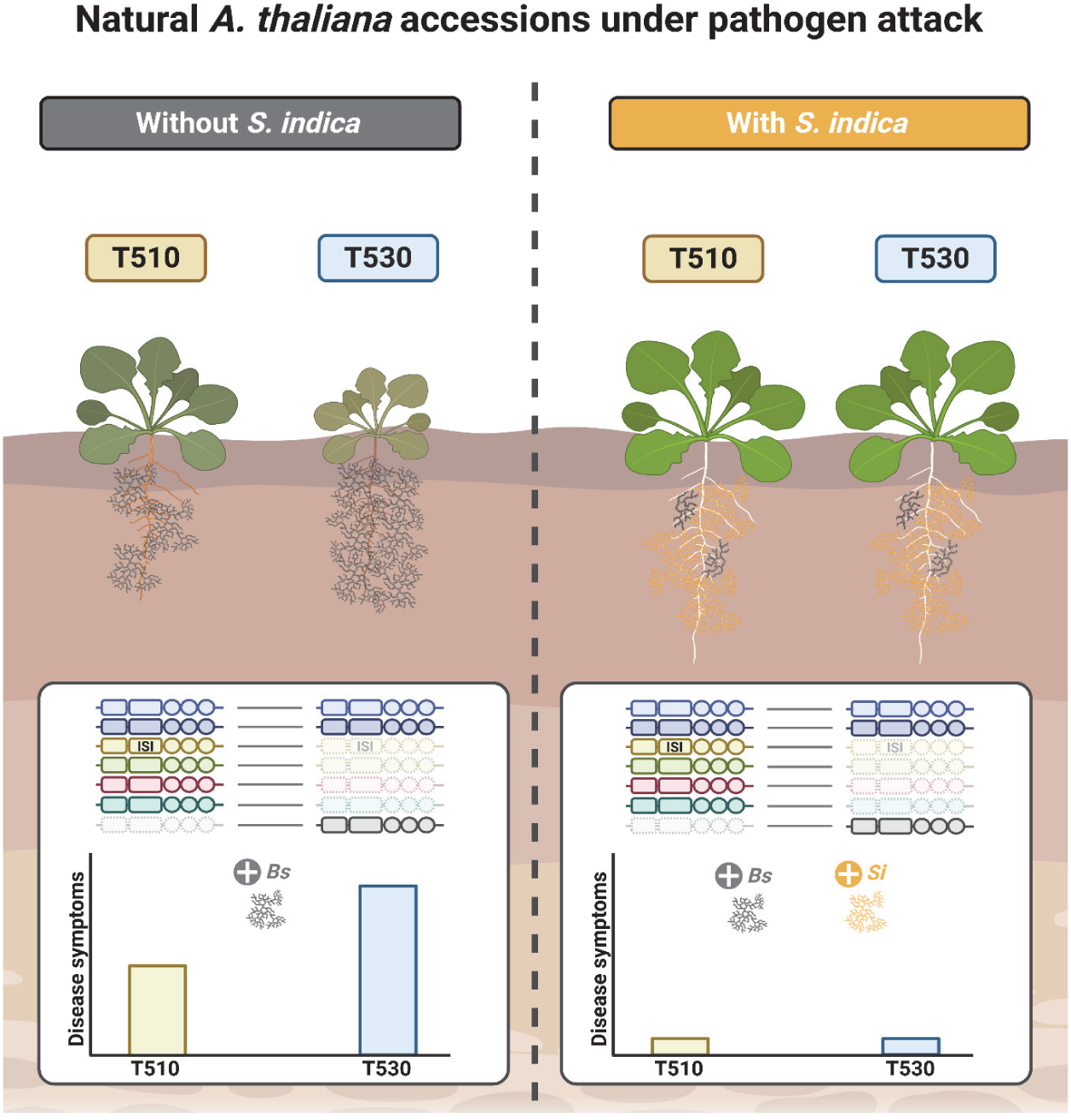
Beneficial symbionts act as ecological stabilizers that mitigate differential pathogen susceptibility within natural plant populations. Presence–absence variation of immune receptor genes generates accession-specific differences in susceptibility to phytopathogens such as *Bipolaris sorokiniana* (*Bs*). In the absence of beneficial microbes such as the root endophyte *Serendipita indica* (*Si*), this immune variation results in differential pathogen colonization and disease severity. When *Si* colonizes the root and rhizosphere, it suppresses *Bs* through extraradical antagonism, greatly reducing pathogen load in both accessions and restoring growth-associated transcriptional programs. By reducing the phenotypic consequences of susceptible immune variants, this microbial protection may buffer pathogen-driven selection pressure, thereby contributing to the persistence of immune-receptor diversity within plant populations.

## Materials and methods

### Plant, fungal and bacterial materials

*Arabidopsis thaliana* ecotype Columbia 0 (Col-0), T510 (NASC ID: N78052), T530 (N78054) and SALK line SALK_034517C (*isi*) were used as plant hosts. Additionally, 45 natural *At* accessions (Table S1) were used which are part of the 1001genomes.org collection (Alonso-Blanco *et al*., 2016). As fungal models *Serendipita indica* (DSM11827, German Collection of Microorganisms and Cell Cultures GmbH, Braunschweig, Germany) and *Bipolaris sorokiniana* (ND90Pr, obtained from S. Zhong, Cereal Disease Laboratory Saint Paul, USA) were used. For bacterial treatments, *Burkholderia glumae* PG1 (Gao *et al*., 2015), obtained from K.-E. Jäger, Heinrich Heine University Düsseldorf, Germany, was used.

### Plant growth conditions and microbial inoculation

Seeds were surface-sterilized twice in 70 and 100 % EtOH for six minutes each and were stratified for two to three days in the dark at 4 °C and subsequently germinated under short-day conditions (8 h light at 22 °C, 16 h at 18 °C dark, 130 μmol m^-2^ s^-1^ light, 60 % humidity) on ½ Murashige-Skoog medium including vitamins (MS, pH 5.7, Duchefa, Haarlem, the Netherlands) supplemented with 1 % (w/v) sucrose and 0.4 % (w/v) Gelrite (Duchefa, Haarlem, the Netherlands). *Si* cultures were propagated on complete medium (CM) (Pham *et al*., 2004) supplemented with 2 % (w/v) glucose and 1.5 % (w/v) agar at 28 °C in darkness for 28 days prior to spore isolation. For mycelial suspensions, liquid CM medium was inoculated with spores and cultured at 28 °C and 120 rpm in the dark. *Bs* propagation was carried out on modified CM at 28 °C in darkness for 14 days prior to spore extraction. Mycelial suspensions (*Si*) and spore suspensions (*Bs*) were prepared as previously described (Hilbert *et al*., 2012; Sarkar *et al*., 2019). Seven-day-old *At* seedlings were transferred to 1/10 Plant Nutrition Gelrite Medium (PNM, pH 5.7, Basiewicz *et al*. (2012)) in 12 × 12 cm square Petri dishes (15-20 seedlings per plate). 1 ml of water containing either 20 mg of *Si* mycelium, 5,000 *Bs* conidia, or a mixture of both was pipetted onto the root and surrounding area. Control plants were inoculated with sterile water (Mock). Prior to harvest, roots were washed thoroughly with sterile water to remove extraradical hyphae. Whole roots 0.5 cm below the shoot were collected. 20-30 roots were pooled per replicate (n = 4) and immediately snap frozen in liquid nitrogen.

Bacterial inoculations were performed using freshly cultured *Burkholderia glumae* PG1. Overnight cultures were washed twice with sterile 10 mM MgCl₂ and adjusted to OD₆₀₀ = 0.0001. Plants received one of four treatments: Mock (110 µl of 0.91 mM MgCl₂), *Bg* (10 µl of bacterial suspension plus 100 µl of milliQ water), *Si* (0.5 mg of *Si* mycelium in 100 µl milliQ water plus 10 µl of 10 mM MgCl₂), or *SiBg* (a mixture of 100 µl milliQ water containing 0.5 mg *Si* mycelium and 10 µl *Bg* suspension) in wells filled with 1 ml ½ MS medium. After three days, roots were harvested for RNA extraction to determine the bacterial load. RNA was isolated using NucleoZOL agent (Marchery-Nagel) according to the manufactureŕs protocol. RT and DNAse treatment were performed using QuantiTect kit (Qiagen). qPCRs were performed using GoTaq qPCR Master Mix (Promega) and primers were chosen to amplify BgNR042931 and the host gene AT4G26410 for normalization (Oligonucleotides listed in Table S11).

### Plant growth conditions and phenotyping in common garden experiment

Plants were grown in two different soil mixes. The nutrient-rich soil (“100 % soil”) consisted of 90 % Einheitserde VM soil (Balster Einheitserdewerk GmbH, Fröndenberg) and 10 % Cologne Agricultural Soil, a nutrient-poor soil (Harbort *et al*., 2020) collected from a natural population site (50.958 N, 6.856 E). The nutrient-poor mix (“30 % soil”) was composed of 70 % sand and 30 % of the nutrient-rich mix. Plants were cultivated in 6×6 cm pots filled with either soil type and placed outdoors under natural environmental conditions in Cologne (50°55’32.0"N 6°56’08.0"E) from September to November. The experiment was conducted in two independent rounds, initiated in 2023 and 2024, respectively. During this period, plants were exposed to ambient light, temperature, and precipitation. Watering was minimal: plants were watered only occasionally, mostly during the early (September) stages of the experiment, when prolonged dryness threatened survival. Rosette growth was monitored by imaging plants from above at regular intervals. Images were analyzed using ImageJ v1.54g (Schneider *et al*., 2012) to measure the rosette diameter. For each plant and timepoint, the longest distance across the rosette was recorded. To account for variation in initial size, rosette diameters were normalized to each plant’s size at day 21 after sowing, and relative growth was computed for all later timepoints.

### Pulse-amplitude modulation (PAM) fluorometry and ion leakage measurement

For PAM fluorometry and ion leakage assays, inoculated *At* plants were used at three to five dpi. Roots were carefully washed to remove fungal hyphae and transferred to 24-well culture plates containing either 2 ml (PAM) or 500 µl (ion leakage) sterile MilliQ water per well. One (PAM) or five (ion leakage) plants per well were used. PAM measurements were taken every 24 h from zero to seven days post transfer (dpt) as described in Dunken *et al*. (2022). Ion leakage was assessed in parallel using a conductivity meter (LAQUAtwin EC-11; Horiba, Newhampton, UK).

### Analysis of screening data

Upon imaging of the 47 screened *At* accessions daily using a PAM imaging fluorometer of the model IMAG-CG (Heinz Walz GmbH, Effeltrich, Germany), two parameters were used for analysis: the maximum quantum yield of photosystem II (FV/FM) and the projected plant area. A plant health index was calculated by multiplying FV/FM by the projected area for each individual plant at each timepoint. To capture overall plant performance over the course of the experiment, health index values were integrated over time to obtain area under the curve (AUC) values for each sample (AUCMock, AUC*Bs*, AUC*SiBs*). From these values, a protection score was calculated for each accession using the formula: (AUC*SiBs* – AUC*Bs*) / AUCMock. Screening was carried out in five experimental rounds with the accession Col-0 included as an internal reference in each of these. Final protection scores were normalized to the Col-0 average within each batch.

### Root length measurements

Root length was measured from plate scans using Fiji (ImageJ) with the NeuronJ plugin by tracing the primary root (Meijering *et al*., 2004; Schneider *et al*., 2012).

### SNP data processing and PCA

Genotype data from 1,135 *At* accessions were obtained from 1001genomes.org. SNPs with minor allele frequency (MAF) ≤ 0.05 were excluded. Linkage disequilibrium (LD) pruning was performed using scikit-allel v 1.3.13 (Miles *et al*., 2024) with a sliding window of 500 SNPs (step size 50, R² < 0.2). PCA was conducted using scikit-learn v1.6.1 (Pedregosa *et al*., 2011) and the first two components were used for visualization.

### RNA extraction for RNA-seq and qPCR

RNA for all data except *Bg* colonization data (see above) was extracted using TRIzol (Invitrogen) and treated with DNase I (Thermo Fisher Scientific) following the manufacturers’ protocols. cDNA was synthesized with the First Strand cDNA Synthesis Kit (Thermo Fisher Scientific). For quantitative real-time PCR, reactions were prepared using the 2× GoTaq qPCR Master Mix (Promega, Mannheim, Germany), supplemented with 500 nM forward and reverse primers and 10–20 ng of cDNA template. Amplification was carried out on a CFX Connect Real-Time System (Bio-Rad, Munich, Germany) using the following program: 95 °C for 3 min; 40 cycles of 95 °C for 15 s, 59 °C for 20 s, and 72 °C for 30 s; followed by melting-curve analysis. Relative gene expression was calculated using the 2^−ΔCT^ (Livak and Schmittgen, 2001). Oligonucleotides are listed in Table S11.

### Transcriptome analyses and genome sequencing

Transcriptome analyses and genome sequencing are described in the Supplemental Materials.

### Statistical analyses

For fungal colonization, plant phenotypes, disease symptoms, and gene expression, data were tested for normality and homogeneity of variances. Depending on the experiment and number of factors, statistical analyses were performed using one-way or two-way ANOVA followed by Tukey’s *post-hoc* test (p < 0.05), or the non-parametric Kruskal–Wallis test when assumptions were not met. If the Kruskal–Wallis test indicated significant differences, pairwise comparisons were performed using Dunn’s test with FDR-adjusted p-values (< 0.05). For direct comparisons between two groups, the Mann–Whitney U test was used. Specific tests applied are indicated in the figure legends.

## Acknowledgements

CS, TT, AY, SK, JDM and AZ acknowledge support from the CRC (Collaborative Research Center) TRR 341 Plant Ecological Genetics funded by the Forschungsgemeinschaft (DFG, German Research Foundation) under Project ID: 456082119. LA, BU, SK, JDM and AZ acknowledge support from the Cluster of Excellence on Plant Sciences (CEPLAS) funded by the DFG under Germany’s Excellence Strategy-EXC 2048/1-Project ID: 390686111. VJR and BU acknowledge support from the Collaborative Research Center MibiNet (CRC 1535), funded by the Deutsche Forschungsgemeinschaft (DFG, German Research Foundation) under Project ID 458090666. VJR also acknowledges the support of Barbara Schulten for sequencing the T510 and T530 accessions. Map data was attained from OpenStreetMap.

## Author contributions

NP, CS, LA, GL and AZ designed the research and conceptualized the manuscript. NP and CS screened natural *At* accessions for *Si*-mediated protection from *Bs* and analyzed the data. NP, AWY, ND, TT and AK characterized T510, T530 and *isi* in-depth and analysed the data. RGC assessed the growth of T510 and T530 in garden experiments. NP, CS and LA analysed the RNA-seq data. VJR and BU sequenced and analysed the genomes of T510 and T530. All authors were involved in editing the paper. AZ, SK, JDM and BU provided funding for the experiments.

## Declaration of generative AI and AI-assisted technologies in the writing process

The authors used ChatGPT (OpenAI, San Francisco, CA) to assist with language refinement and clarity improvements. Perplexity.ai (Perplexity AI Inc., San Francisco, CA) was used to support fact-checking during the manuscript preparation. All content was written, verified, and approved by the authors without reliance on AI for original scientific interpretation or conclusions.

## Data availability

All data supporting the findings of this study are available within the article and supporting information and can be downloaded from the ARC repository (doi ARC). The genomes of T510 (ERS27376026, BioSample SAMEA120578005) and T530 (ERS27376027, BioSample SAMEA120578006) have been uploaded to ENA (European Nucleotide Archive). RNA-seq data discussed in this study was deposited in NCBI’s Gene Expression Omnibus (Edgar *et al*., 2002) and are accessible through GEO Series accession number GSE307677 (https://www.ncbi.nlm.nih.gov/geo/query/acc.cgi?acc=GSE307677).

