## Supplemental Methods and Figures for "A fungal root endophyte functionally complements host immunity and mitigates natural immune variation in Arabidopsis"

#### Transcriptomic data analysis

Samples were prepared of accession T510, T530 and Col-0. Per Sample, 20-30 whole roots have been harvested at two timepoints (3 and 6 dpi) under 4 conditions: Mock (water), *Si*, *Bs* and *SiBs*. Each sample was prepared with 3 replicates. mRNA-Seq libraries were prepared using the Illumina TruSeq stranded according to the manufacturer's instructions (Illumina Inc., San Diego, CA, USA). Sequencing was performed on the NovaSeq6000 platform at the Cologne Center for Genomics (CCG). Samples submitted for sequencing contained 2 µg of total RNA in 20 µl $ddH_2O$, with RNA quality indicators $OD_{260/280}$ between 1.8 and 2.1 and $OD_{260/230}$ >1.5. For each of the three biological replicates, approximately 100 million paired-end reads of 100 base pairs in length were generated. Raw paired-end RNA-seq reads were processed with fastp v0.20.0 (Chen *et al*., 2018a) to remove adapter sequences and low-quality bases, discarding reads shorter than 98 bp. Quality control of trimmed reads was conducted using FastQC v0.11.9 (Andrews, 2010) and summarized across samples with MultiQC v1.19 (Ewels *et al*., 2016). Reads were aligned to the *At*, *Si* and *Bs* reference genomes TAIR10 (Lamesch *et al*., 2012), DSM 11827 (Zuccaro *et al*., 2011) and ND90Pr v1.0 (Condon *et al*., 2013; Ohm *et al*., 2012) using STAR v2.7.10a (Dobin *et al*., 2013). Mapping statistics for each genome are shown in Table S2 (*At*), S3 (*Si*) and S4 (*Bs*). Gene-level counts were obtained by aligning reads to the *Arabidopsis thaliana* TAIR10 reference genome (Ensembl Plants release 58) using the corresponding gene annotation, and imported into R for differential expression analysis with DESeq2. (Love *et al*., 2014). Genes <10 counts in all samples of the smallest condition group were excluded. Normalization and dispersion estimation were performed using the DESeq() function. For visualization, variance-stabilizing and regularized log transformations were computed. Differential expression was assessed separately for each genotype at 3 dpi, using models appropriate to the number of available treatment groups. $Log_2$ fold changes were shrunk using the apeglm method (Zhu *et al*., 2019). Genes with an adjusted p-value ≤ 0.05 and $|log_2FC| \geq 1$ were considered differentially expressed. Gene ontology (GO) term enrichment analysis was performed using the enrichGO function of the clusterProfiler package with "org.At.tair.db" as an organism database (Yu *et al*., 2012) to identify biological processes overrepresented among DEGs relative to the Col-0 genome background. Enriched GO terms were grouped into manually curated functional categories.

#### Genome sequencing, structural variation and genome annotation

Plants of accessions T510 and T530 were cultivated under controlled greenhouse conditions. For each accession, 1.5 g of fresh leaf tissue was used for DNA extraction as previously described (Ziegler *et al*., 2025). In brief, the NucleoBond HMW DNA Extraction Kit (Macherey-Nagel, Germany) was used following the manufacturer's instructions. DNA quality and quantity were assessed using a NanoDrop spectrophotometer (Thermo Fisher Scientific, USA) and the Qubit dsDNA BR Assay Kit (Thermo Fisher Scientific, USA). 9000 ng high-quality DNA per sample was size-selected using the PacBio SRE XL Size Selection Kit (Pacific Biosciences, USA). Library preparation was performed using the Oxford Nanopore SQK-LSK114 XL Ligation Sequencing Kit (Oxford Nanopore Technologies, UK) according to the manufacturer's protocol. Sequencing was conducted on Oxford Nanopore R10.4.1 FLO-PRO114M flow cells using a PromethION platform (Oxford Nanopore Technologies, UK).

In total, approximately 89 Gb were generated for T510 and 105 Gb for T530. Basecalling was performed using Dorado version 0.9.1 (Oxford Nanopore Technologies, UK) with the super-accurate model (sup@v5.0.0, 5mC_5hmC). The resulting reads were filtered with SAMtools v1.20 (Danecek et al., 2021), applying a minimum Q-score of 15 for T510 and 20 for T530, using the command-line options view -e "[qs] >= 15" for T510 and view -e "[qs] >= 20" for T530. Genome assembly was carried out using hifiasm version 0.24 (Cheng *et al*., 2021). For each accession, reads with lengths between 15 kb and 50 kb were provided via the --ont option, while reads longer than 50 kb were included via the --ul option.

To identify chromosomes, the resulting assemblies were compared to the Arabidopsis thaliana Col-0 reference genome (TAIR10; Berardini *et al*. (2015)) using D-GENIES (Cabanettes and Klopp, 2018). Chromosomal sequences were extracted with seqtk version 1.4 using the subseq command. The chromosome-scale assemblies were subsequently polished with NextPolish (Hu *et al*., 2020) using the --lgs_option. A reference-free error correction step was then performed with Inspector version 1.3 (Chen *et al*., 2021) using the --datatype nano-raw option. Assembly completeness was evaluated with Compleasm version 0.2.7 (Huang and Li, 2023) using the brassicales_odb12 model. Assembly quality metrics were generated with Merqury version 1.3 (Rhie *et al*. (2020); parameter k = 23; Table S8) and Craq version 1.0.9 (Li *et al*., 2023).

Genome annotation was performed using three complementary strategies. First, the TAIR10 annotation was lifted to the T510 and T530 genomes using Lifton version 1.0.4 (Chao *et al*., 2025; Shumate and Salzberg, 2021). Available RNA-seq data for T510 and T530 were quality-trimmed with fastp version 0.24 (Chen *et al*., 2018a) and subsequently mapped using HISAT2 version 2.2.1 (Kim *et al*., 2019) with the parameters --dta and -x. The resulting BAM files were merged with SAMtools version 1.20 (Danecek *et al*., 2021) using the merge command. The lifted TAIR10 annotation and the HISAT2 alignment files served as input for StringTie version 3.0.0 (Pertea *et al*., 2015) with the options -G, -c 5, -j 3, and -f 0.1. As an additional ab initio approach, Helixer version 0.3.3 (Holst *et al*., 2023; Stiehler *et al*., 2021) was applied using the land-plant model (v0.3_a_0080) with default parameters. The annotations generated by StringTie and Helixer were then combined using Mikado version 2.3.3 (Venturini *et al*., 2018). To ensure that only valid splice junctions were considered, Portcullis version 1.2.4 (Mapleson *et al*., 2018) was run beforehand on the RNA-seq alignments using the portcullis full command. Finally, the resulting annotation file was filtered for isoforms with AGAT version 1.4.0 using the agat_sp_keep_longest_isoform script (Dainat, 2022). Completeness of the predicted proteomes was assessed using Compleasm version 0.2.7 (Huang and Li, 2023) with the brassicales_odb12 lineage dataset.

Structural variation and synteny analyses were performed using SyRI version 1.7.0 (Goel *et al*., 2019) for whole-genome structural visualization. For both macro- and microsynteny analyses, the Mikado-derived CDS and GFF files, together with the *A. thaliana* Araport11 reference (Cheng *et al*., 2017), were used. Visualization was carried out with JCVI version 1.5.6 (Tang *et al*., 2024). During orthology inference with jcvi.compara.catalog ortholog, the --cscore parameter was set to 0.95. In subsequent analyses using jcvi.compara.synteny screen, the --minspan parameter was set to 10. The resulting anchor file used for macrosynteny visualization was then filtered to retain only the best reciprocal hits, thereby removing multiple mappings. Finally, jcvi.compara.synteny mcscan was executed with the parameter --iter=1 Subsequently, synteny was inspected and gene models improved manually.

For the presence–absence analysis of the candidate genes, annotation-free approaches were applied. In total, 58 candidate genes were analyzed for strain T510 and 48 candidate genes for strain T530. First, the complete nucleotide sequences of the candidate genes, taken from the updated *At* annotation (Araport11) were aligned against their corresponding reference genomes using BLASTn version 2.15 (Altschul *et al*., 1990; Camacho *et al*., 2009). Subsequently, the protein sequences of the candidates were aligned to the genomes with Miniprot version 0.14 (Li, 2023) using the options -I, -u, and --gff.

Comparison of both alignment results provided clear evidence regarding the presence or absence of each gene in the T510 and T530 genomes. Candidate protein-coding genes with Miniprot alignments were inspected in the genome annotation file of the respective genome to identify overlapping or nearby genes. These genes were then extracted and further examined by manual BLASTp searches against the TAIR10 protein database.

### Supplemental Figures

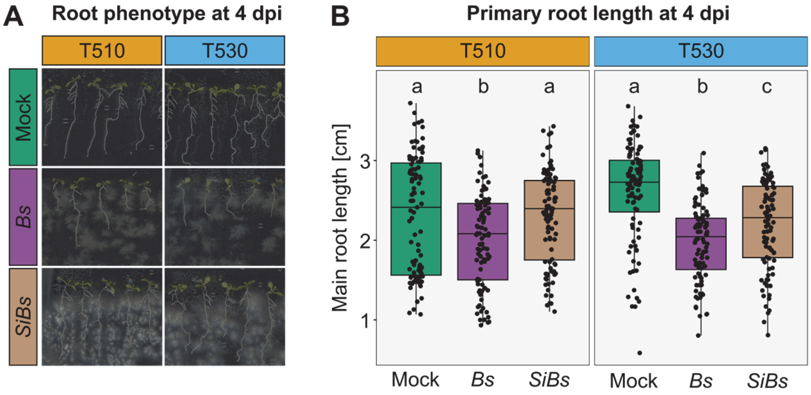


**Figure S1: Effect of *Bs* and *SiBs* on *At* root morphology. (A) Representative root phenotype at 4 dpi.** Seven-day-old T510 and T530 seedlings were treated with *Bs*, *SiBs*, or water (Mock) on 1/10 PNM medium. Pictures were taken every day to document the root phenotype. **(B) Quantification of primary root length at 4 dpi (n = 96).** Statistical significance was assessed using ANOVA with Tukey's *post-hoc* test. Different letters indicate significant differences of treatments within genotypes ($p < 0.05$).

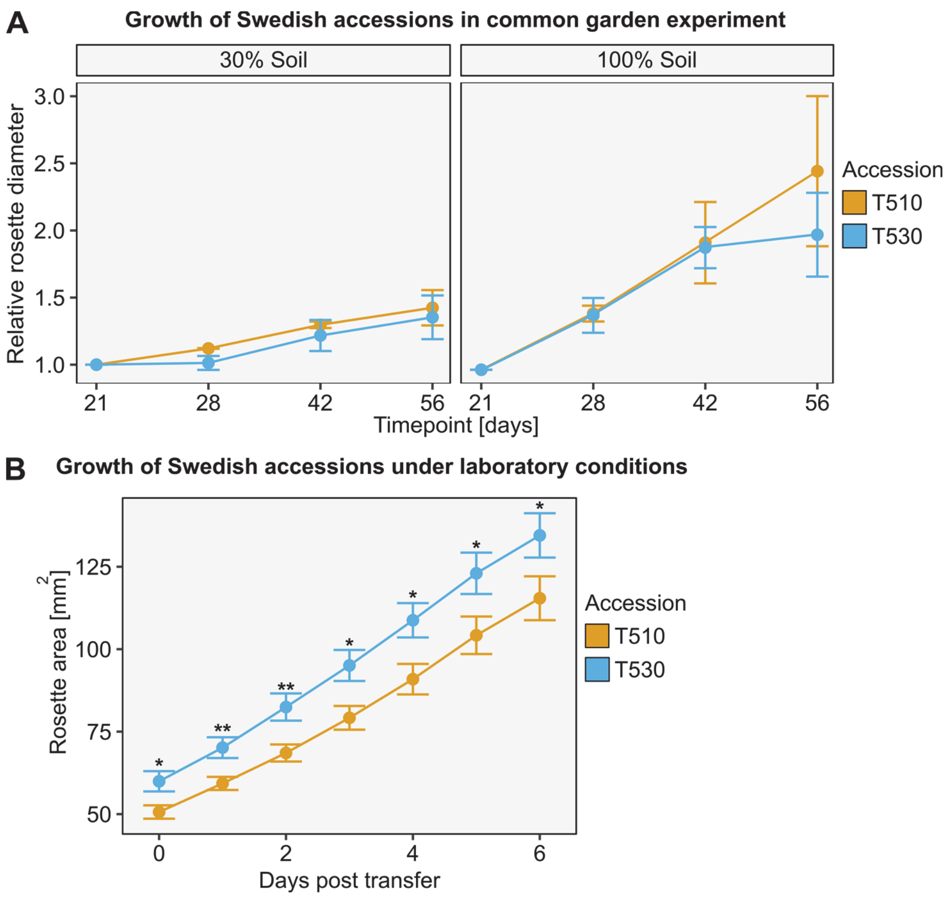


**Figure S2: Growth of Swedish accessions under different conditions. (A) Garden experiment with T510 and T530 grown under low (left panel) and high (right panel) nutrient conditions.** Rosette diameter over time is shown for plants grown in nutrient-poor (30 % soil) or nutrient-rich (100 %) substrates. The nutrient-rich mix consisted of 90 % Einheitserde VM mixed with 10 % Cologne Agricultural Soil, a nutrient-poor soil. The nutrient-poor mix comprised 70 % sand and 30 % of the nutrient-rich substrate. Data from two independent experimental rounds were combined (n = 6). Rosette diameter at each timepoint was normalized to the value at day 21 after sowing to visualize relative growth. Dots indicate the mean relative diameter; error bars represent the standard error of the mean (SEM). **(B) Growth of T510 and T530 under laboratory conditions.** Seedlings were grown under the same conditions as described for the microbial inoculations followed by PAM measurements. In short, seedlings were germinated on ½ MS. After seven days, the plants were transferred to 1/10 PNM Gelrite medium and inoculated with water. After four days, the plants were transferred to 24-well plates containing 1/10 PNM. Leaf surface area was quantified daily after transfer (n = 24). Statistical significance was assessed after testing for normality and homogeneity of variances. Depending on the results, either a Student's t-test or a paired Wilcoxon rank sum test was applied. Dots indicate the mean rosette area; error bars represent the SEM. Asterisks indicate p-values (* $p < 0.05$, ** $p < 0.01$, *** $p < 0.001$).

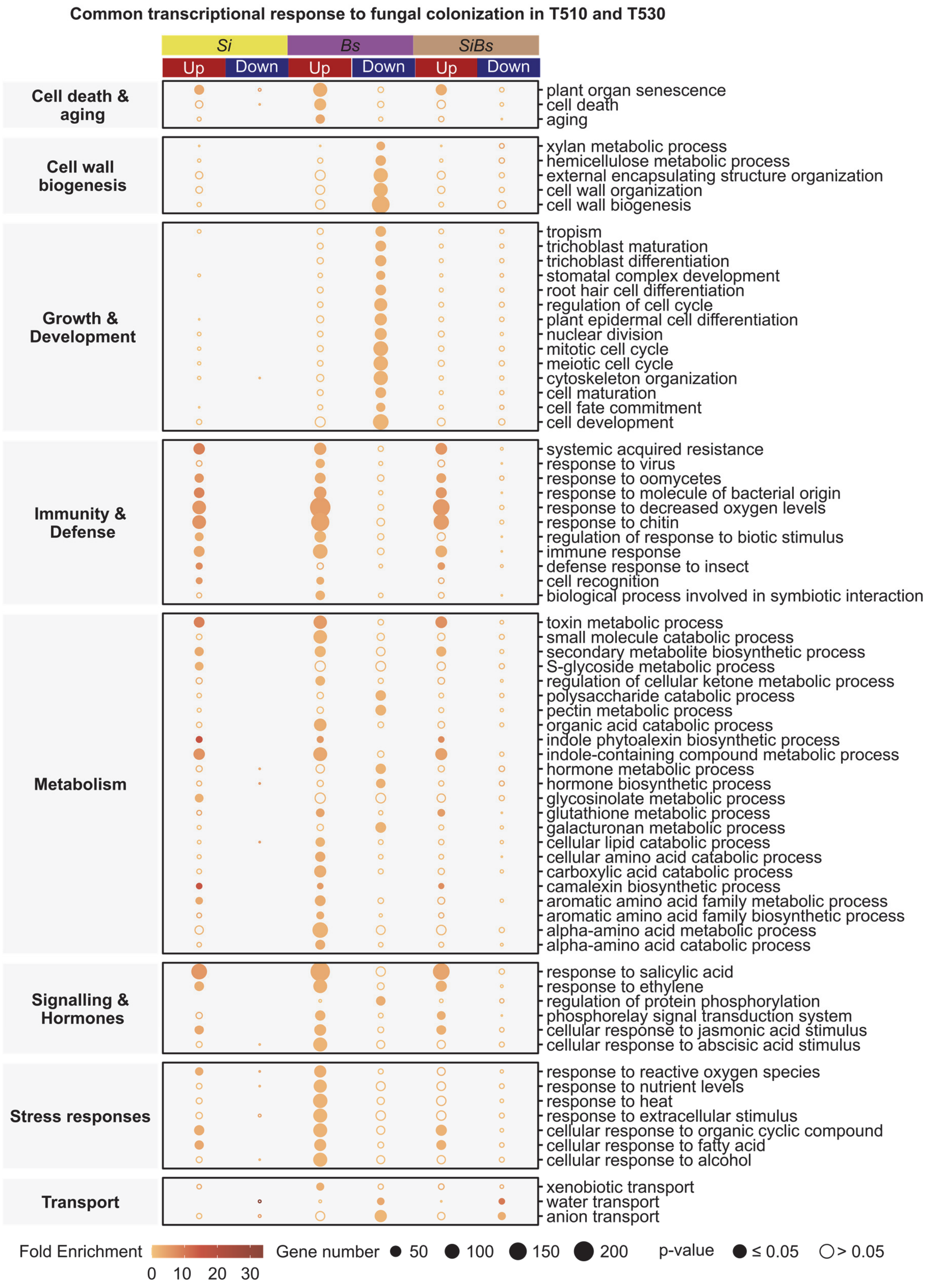


**Figure S3: GO term enrichment analysis of common transcriptional response to fungal colonization in T510 and T530.** GO terms enriched among DEGs responsive ($log_2FC \geq 1$ (up) or $log_2FC \leq -1$ (down) and p.adj ≤ 0.05) to *Bs*, *Si* or *SiBs* in both T510 or T530 (n =3). Dot color indicates fold enrichment, fill denotes statistical significance, and size reflects the number of genes associated with each GO term.

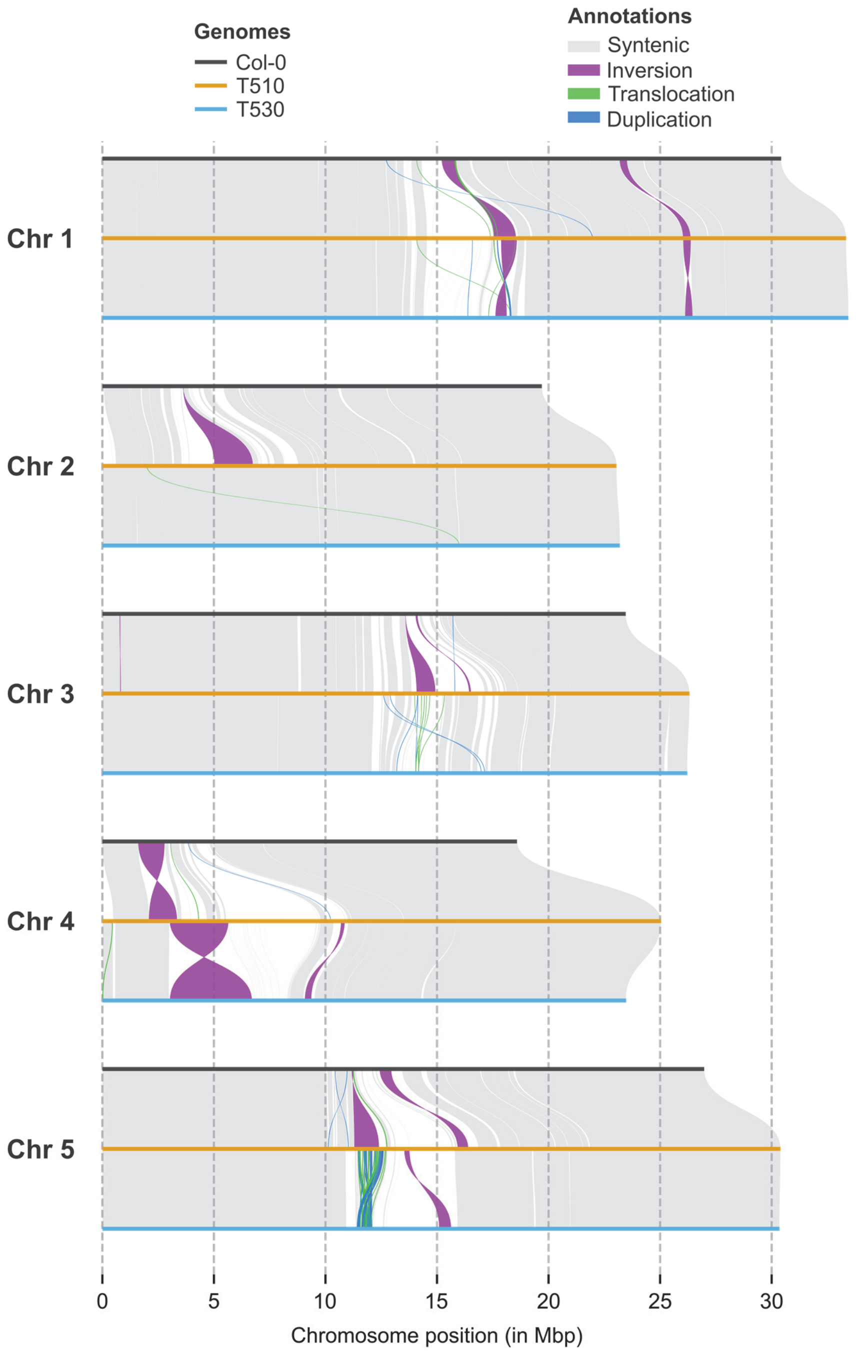


**Figure S4. Structural differences between the genomes of T510, T530, and Col-0.** Genome alignments reveal a high degree of synteny between accessions T510 and T530 relative to the *At* Col-0 reference genome. Across all chromosomes, smaller inversions (depicted in purple) are visible. The most prominent structural variation is a larger inversion on chromosome 4 observed between T510 and T530.
